# Microplastics influence size-selected zebrafish behaviour

**DOI:** 10.64898/2026.03.06.710120

**Authors:** Daniel E. Sadler, Stephan N. van Dijk, Silva Uusi-Heikkilä

## Abstract

Plastic pollution represents a major contemporary threat to aquatic ecosystems, with well-documented consequences for organismal performance and fitness across numerous taxa, including fishes. Importantly, plastic-derived stress does not occur in isolation, but interacts with other anthropogenic pressures such as size-selective harvesting, which can impose strong directional selection on life-history and behavioural traits. In this study, we exposed three experimentally evolved selection lines: large-harvested, small-harvested, and randomly harvested to microplastic contamination and quantified effects on growth and behaviour over a 14-day period. Microplastic exposure reduced boldness and exploratory activity while simultaneously increasing feeding probability and feeding frequency. Prior size-selective harvesting influenced only exploratory behaviour, suggesting that most behavioural responses to microplastics are robust to previous evolutionary history. We detected no effect of microplastics on growth, potentially due to compensatory increases in feeding behaviour. Collectively, these findings demonstrate that microplastic exposure alters key behavioural traits across genetically divergent fish lines and contribute to a broader understanding of how multiple anthropogenic stressors may interact to shape population dynamics in rapidly changing environments.

## 1. Introduction

Aquatic ecosystems are increasingly exposed to a suite of human-induced stressors including shifts in water temperature (Mittelbach et al., 2007; Parmesan and Yohe, 2003), pH (Caldeira and Wickett, 2003), hypoxia (Feely et al., 2018), and habitat destruction (Malmqvist and Rundle, 2002; Rossi, 2013). At the same time, pollutants have become pervasive in aquatic environments (Andrady, 2011; Young et al., 2016), with microplastics emerging as a particularly widespread and persistent contaminant (Vivekanand et al., 2021; Xu et al., 2020). Microplastics are defined as plastic particles smaller than 5 mm (Rochman, 2018), and originate either as primarily produced particles released through industrial processes and wastewater or as secondarily fragments produced by the degradation of larger plastic debris (Song et al., 2017). Microplastics have an extensive distribution, long environmental persistence and high likelihood of ingestion by aquatic organisms, posing significant ecological risks (Cole et al., 2013; Ivleva et al., 2017). Beyond individual level effects, microplastics can be transferred from prey to predator, enabling bioaccumulation through entire food webs (Carbery et al., 2018; Setälä et al., 2014). A key trait of microplastics is that they can further fragment into nanoplastics (>1000nm; Gigault et al., 2018; Koelmans et al., 2015), which may be even more biologically harmful than microplastics due to their enhanced ability to penetrate tissues (Aransiola et al., 2025), disrupt cellular processes (Aransiola et al., 2025; Jeong et al., 2024), accumulate more readily (Rist et al., 2017), and trigger immune responses (Sadler et al., 2019).

In fish, exposure has been linked to reductions in feeding and growth rates (Barría et al., 2020), impaired locomotion (Mattsson et al., 2017; Pitt et al., 2018), genotoxic effects (Menezes et al., 2024), decreased reproductive success (Wang et al., 2019) and reduced survival (Manabe et al., 2011). Microplastics have been detected in fish digestive tract, gills, and liver (Ferreira et al., 2016) indicating that they can influence multiple physiological systems simultaneously. Despite this, most existing studies focus on isolated traits within an ecotoxicological framework, and comparatively few study their effects by integrating morphology, life-history and behavioural traits. Such trait complexes are often interdependent and may provide mechanistic insight into how microplastics disrupt organismal performance.

Animals operate under a finite energy budget; therefore, resources must be allocated among essential functions such as growth, reproduction, activity, maintenance (e.g., immune activity and cellular repair), and storage (e.g., lipid reserves). Life history strategies differ in how this allocation is structured: individuals with fast life history strategies typically grow faster, mature earlier at smaller sizes, and have shorter lifespans compared to slow strategy individuals (Healy et al., 2019). These contrasting allocation pathways suggest microplastic exposure may disproportionately affect life-history types. If microplastics divert energy away from somatic growth or other key functions, fast strategy individuals may fail to mature and reproduce at an optimal age (Shang et al., 2021), with potential consequences for population dynamics and broader ecosystem functioning.

Beyond life history and physiological processes, behaviour is a critical determinant of individual fitness (Smith and Blumstein, 2008) and often represents the first line of defence organisms can deploy to buffer against environmental stressors (Wong and Candolin, 2015). Microplastics have been shown to alter behaviour in fishes (Mattsson et al., 2017; Pitt et al., 2018; Salerno et al., 2021), including traits associated with animal personality, defined as consistent individual differences across time in response to different stimuli (Boissy, 1995; Brown et al., 2011; Sih et al., 2004). Indeed, recent evidence suggests that microplastic exposure can modify personality traits in fish (Chen et al., 2022). Among these traits, boldness and exploration are of particular importance. Bold individuals often forage more actively (Magnhagen and Staffan, 2003) and achieve higher reproductive success (Nakayama et al., 2017), while exploratory behaviour influences resource acquisition and social interactions (Conrad et al., 2011). Empirical studies demonstrate that microplastics can disrupt these behaviours. For example, microplastic exposure reduced swim speed in black rockfish (*Sebastes schlegelii;* Yin et al., 2019), induced anxiety like responses in zebrafish (*Danio rerio*; Félix et al., 2023), and led to hyperactive swimming (Chen et al., 2020). Microplastics may also alter feeding behaviour, with exposed individuals spending nearly twice as long feeding (Mattsson et al., 2017). Such behavioural changes have the potential to reshape energy allocation pathways and consequently, influence key fitness components including growth and reproduction.

The effects of plastic pollutants do not occur in isolation, rather, they act concomitantly with other environmental stressors, potentially producing antagonistic or synergistic interactions. A major stressor in aquatic systems is overfishing, which is frequently size-selective and can drive evolutionary shifts towards faster growth rate, earlier maturation and altered physiological traits (Olsen et al., 2004; Reid et al., 2023; Uusi-Heikkilä et al., 2015; van Wijk et al., 2013). Size-selective harvesting also influences behaviour, including feeding activity, general activity levels, exploration, and boldness (Sadler et al., 2024; Sbragaglia et al., 2019; Uusi-Heikkilä et al., 2015). Consequently, it is plausible that size-selective harvesting and pollutant exposure interact to shape behavioural responses in fish. Size-selection can favour particular personality types, such as increased boldness (Uusi-Heikkilä et al., 2015), while pollutants like microplastics can simultaneously affect the same behavioural traits (Chen et al., 2022). Indeed, recent work demonstrates that pollutants and size-selection can interact to favour slow life-history strategies (Uusi-Heikkilä et al., 2024), which may in turn influence behavioural phenotypes. Understanding how fisheries-induced selection and pollutant exposure jointly affect fish biology is particularly important within a fisheries context.

Here, we used an experimental model system to study how small microplastic exposure (500nm particles) affected fish condition factor (energy storage), growth, and behaviour across three different selection lines. Two of the lines were exposed to directional selection for small and large body size and one was selected randomly with respect to body size (Uusi-Heikkilä et al., 2015). Selection favouring small body size typically favours fast life-history strategy with fast juvenile growth and early maturation (Heino et al., 2015; Jørgensen et al., 2007; Uusi-Heikkilä et al., 2015), and small-selected fish have been shown to be less resilient towards other aquatic pollutants (Uusi-Heikkilä et al. 2024), therefore, we hypothesised that fish from different selection lines would respond differently to microplastic exposure. Consequently, we predicted that exposure to microplastics may alter the energy allocation pathways by (1) affecting behavioural traits, such as activity and feeding rate, and consequently alter condition factor and growth rate.

## 2. Material and methods

### 2.1. Study system

Wild-caught zebrafish originating from West Bengal, India were subjected to five generations of size-selective harvesting under three distinct selection regimes (1) small-selected, in which 75% of the largest individuals were removed (typical fisheries selection against large body size); (2) large-selected, in which 75% of the smallest individuals were removed; and random-selected, in which 75% of individuals were removed irrespective of body size (a control line without directional selection). Each selection line consisted of two replicates of 450 individuals each. Full details of the experimental design are provided in Uusi-Heikkilä et al. (2015). All procedures were approved by the Finnish Project Authorisation Board (Licence no. ESAVI/24,875/2018) and all experiments adhered to the ARRIVE guidelines (Percie du Sert et al., 2020).

### 2.2. Microplastics

We used carboxylate-modified polystyrene beads with fluorescent red colouring (505 nm excitation, 550 emission; Sigma-Aldrich, UK) which had an average particle size of 500 nm in a 2.5% w/v aqueous suspension. We diluted the suspension to 1.5 g/L with molecular grade deionized water as a working solution. This working solution was vortexed each time before adding to the experimental tanks, creating a concentration of 1 mg/L per tank.

### 2.3. Experimental setup

Prior to the microplastic exposure experiment adult zebrafish were kept at 28°C with a 14:10 (L:D) light cycle and fed ad libitum with a mixture of dry food (TetraMin XL) and live *Artemia salina*. Fish were placed individually in ten 1L tanks per selection line per treatment (microplastic exposure and control). Although fish were kept individually, tanks were placed in close proximity, so they were able to see each other to prevent social anxiety (Shams et al., 2017). An air circulation system was established among the small experimental tanks. Fish were acclimated in their tanks for one week, after which they were exposed to microplastics directly added to the water. Exposure lasted for 14 days during which microplastics were added to the tank twice at the same concentrations due to water exchanges each week to prevent nitrate and waste buildup.

### 2.4. Growth rate and condition factor

Standard length (SL) and wet mass (WM) of all fish were recorded before and after the experimental period by placing individuals under anaesthesia (2-phenoloxyethanol, 1.5% concentration). To determine SL, fish were photographed (against millimetre paper for scale) using a Canon EOS 90D DSLR Camera affixed with a Sigma 105 mm DG Macro HSM lens. Images were measured using ImageJ (Schneider et al., 2012). WM was measured using an analytical balance (Mettler AE240). Specific growth rate was calculated for WM and SL (ln final length (or weight) − ln initial length (or weight)/days × 100). Condition factor (K) was calculated according to Fulton (1911): K = 100 × SL (mm)/WM (g)^3^.

### 2.5. Behavioural trials

We measured the effect of microplastic exposure on exploration, boldness, activity, and feeding behaviour. Fish were starved 24 h before the trials to prevent any effect of feeding and digestion which could have had confounding effects on the feeding behaviour assays. Behavioural trials occurred in a glass tank (30 L) divided into two sections by an opaque plastic sheet. One compartment was darkened and acted as a refuge, while the other compartment contained stones and novel objects (coloured tiles) and thus acted as an area for exploration (see Sadler et al., 2024 for full details). Fish were acclimated in the refuge area for 10 min before the divider was removed and then fish were allowed to explore the novel environment for 20 min. Exploration was measured as time spent exploring a novel environment (Conrad et al., 2011; Le Roy et al., 2021), boldness was measured as the latency of emergence from a refuge (Frost et al., 2006; Krause et al., 1998), and activity was defined as total number of emergences from a refuge.

Feeding behaviour was recorded for five min by adding approximately six mg of flake food (TetraMin XL) and measuring the frequency at which fish took food from the water surface and the time taken to start feeding. After each trial, excess food was removed, and water replaced in the tank. Behaviour was filmed throughout the duration with an overhead view and a side view using a GoPro 7 Silver and a Canon EOS 90D. All trials were performed within an isolated, temperature-controlled room to prevent disturbance. Behaviour videos were viewed to measure exploration time (s), number of emergences, latency of emergence (s), feeding frequency (number of feeding events), probability to feed and latency to feed (s).

### 2.6. Statistical analysis

All statistics were conducted using R v. 4.5.1. We used linear mixed effect models (LMM) and generalised mixed effect models (GLMM) to analyse the effect of microplastics, selection line and their interaction on growth rate (WM and SL) and behavioural traits (activity, boldness, exploration, and feeding behaviour). Selection line replication was set as a random effect.

For the behavioural data, all trials were repeated (i.e. 2 measures for each individual), and trial repeat was included as a random factor in the models to account for repeatability. Additionally, when directly tested, fish ID had no significant effect on any behavioural trait (p>0.05). We used *lmer* and *glmer* within the lme4 package (Bates et al., 2015) and the *lmertest* function within the lmerTest package (Kuznetsova et al., 2017). For data that did not fit the assumptions of a LMM (non-normality and heterogeneity) we used a GLMM, for feeding probability *family = binomial* and number of feeds *family = negative binomial*. Post hoc pairwise comparisons of significant interactions were made using Tukey contrasts with *emmeans* function within the emmeans package (Lenth, et al., 2018).

## 3. Results

### 3.1. Behaviour

Fish exposed to microplastics were less bold than the control fish (LMM: F_1,110_=7.503, p<0.01), while the selection line had no significant effect on boldness (Figure 1a; Table S1). Activity was not influenced by exposure to microplastic or by selection line (Figure 1b; Table S1). There was a significant interaction between microplastic exposure and selection line in explorative behaviour (Figure 1c; Table S1). Small-(emmeans contrast: 0.630, p<0.001) and random-selected lines (emmeans contrast: 1.154, p<0.001) exposed to microplastics were less explorative than the control fish, but large-selected (emmeans contrast: -0.131, p<0.05) and control fish did not differ in their explorative behaviour.

**Figure 1.**
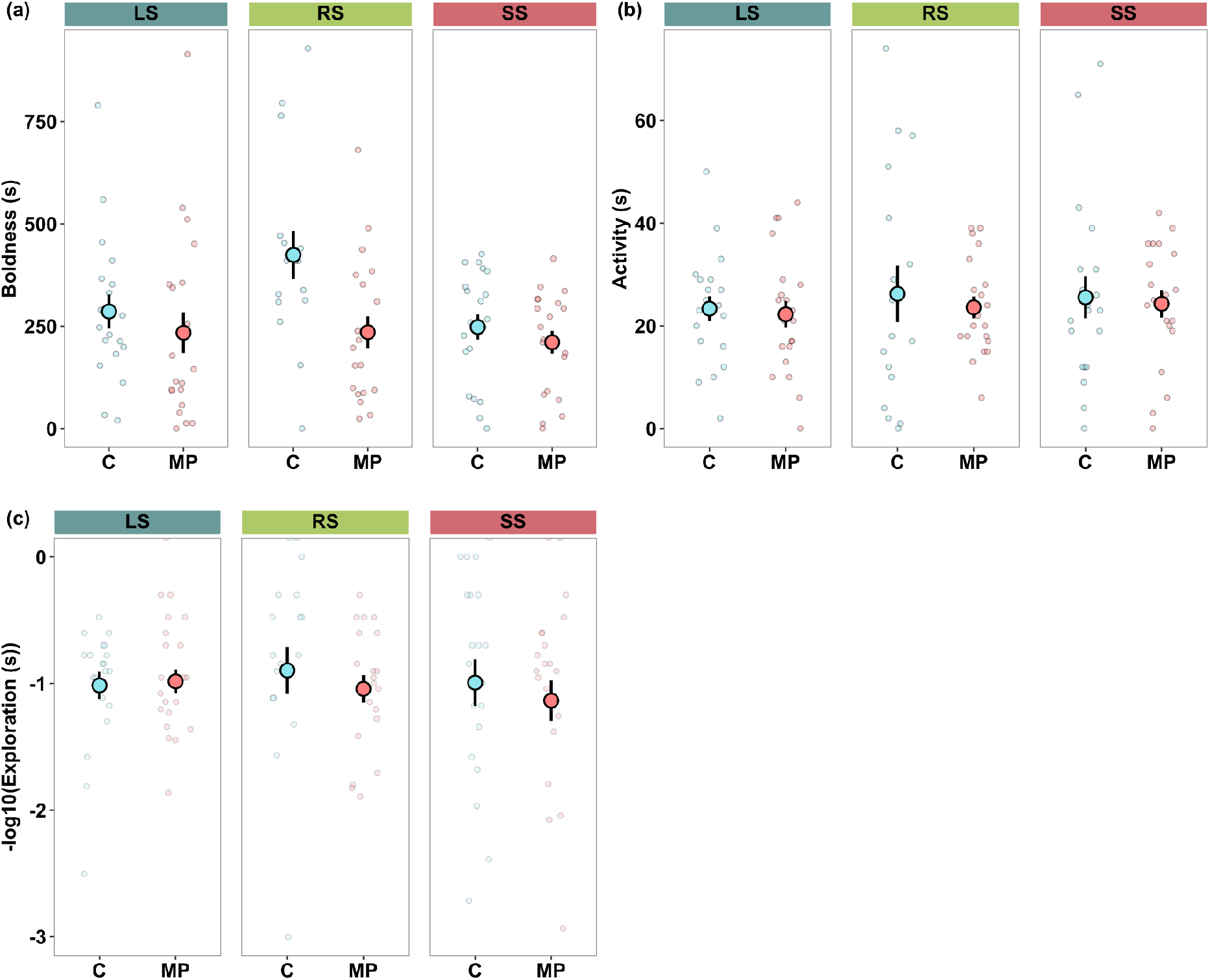
Differences in (a) boldness, (b) activity, and (c) exploration (transformed with - log10 to allow for visualisation) between control fish (C) and fish exposed to microplastics (MP) across selection lines. Large-selected (LS), random-selected (RS), and small-selected (SS). Large dots indicate mean ± CI. Small dots indicate individual measurements.

Feeding latency was not affected by microplastic exposure or by selection line, (Figure 2a; Table S1). The number of feeding events was greater in fish exposed to microplastics (GLMM: z=2.04, p<0.05) but did not differ between selection lines (Figure 2b; Table S1). Microplastic exposure increased probability to feed (GLMM: F_1,110_ = 3.394, p<0.001) but this did not differ among the selection lines (Figure 2c; Table S1).

**Figure 2.**
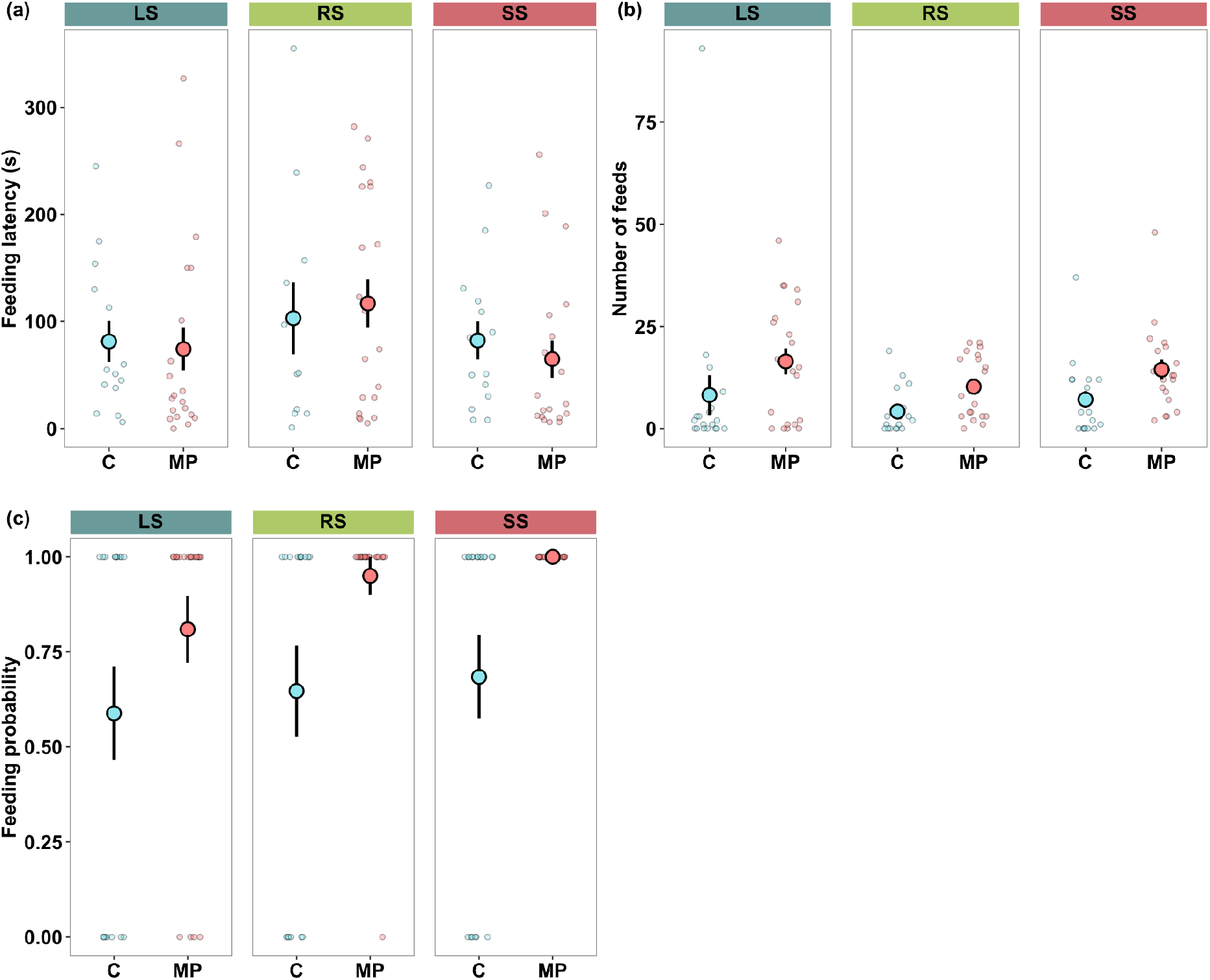
Differences in (a) feeding latency, (b) number of feeds, and (c) feeding probability between control (C) and microplastic exposed (MP) fish across the selection lines. Large-selected (LS), random-selected (RS), and small-selected (SS). Large dots indicate mean ± CI. Small dots indicate individual replicates.

### 3.2. Growth rate and condition factor

Microplastic exposure had no significant effect on either condition factor (Figure 3a; Table S1) or growth rate (Figures 3b, S1; Table S1). Growth rate and body condition did not significantly differ between selection line replicates (Figure 3; Table S1).

**Figure 3.**
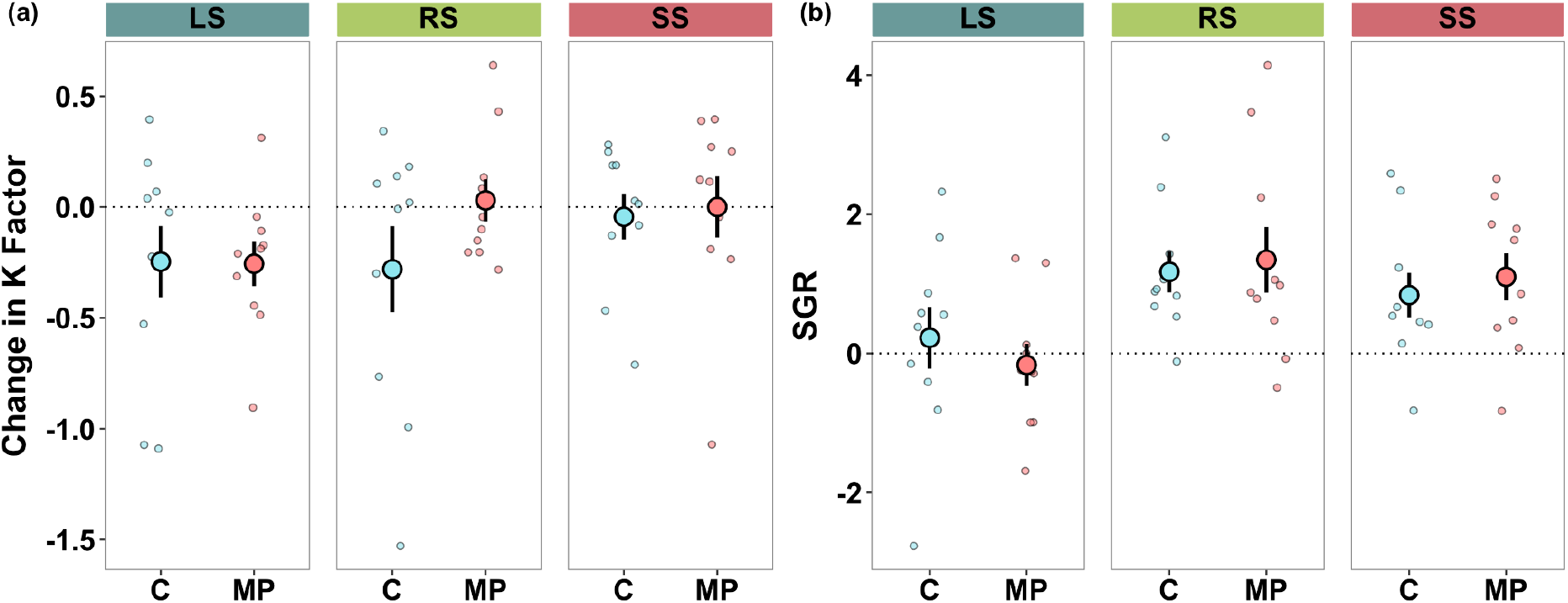
Differences in (a) condition factor, and (b) specific growth rate (SGR) in length between control (C) and microplastic exposed fish (MP) across the selection lines. Large-selected (LS), random-selected (RS), and small-selected (SS). Condition factor (K) was calculated at the start and end of the experiment, with the change shown here (end - start). Large dots indicate mean ± CI. Small dots indicate individual replicates.

## 4. Discussion

Microplastics are ubiquitous across aquatic environments (Rochman, 2018) and have been shown to have profound effects on physiology and behaviour in fishes (Sequeira et al., 2020; Wang et al., 2020). We showed that zebrafish exhibited distinct behavioural responses when exposed to microplastics. Fish exposed to microplastics were less bold and explorative compared to the control fish. At the same time, they showed increased feeding probability and a higher number of feeding attempts. Increased feeding efficiency could be a strategy to compensate for the potential negative effects of microplastics on growth and condition factor. Differences among the selection lines were generally absent, implying that prior size-selective harvesting did not strongly modulate responses to microplastics. The only exception was exploration behaviour, where small-selected fish displayed reduced exploration. Overall, these findings demonstrate that short-term microplastic exposure affects behavioural traits more readily than growth, and that prior size-selective harvesting plays a limited role in shaping these responses.

Despite extensive evidence that microplastics affect growth rate in fish primarily through impaired feeding and blocked guts (Barría et al., 2020; de Sá et al., 2018; Qiao et al., 2019), we found no effect of microplastic exposure on growth rate. This may be due to the duration of the experiment (14 days). Many previous studies have exposed fish to microplastics over a longer period of time, for example Naidoo and Glassom (2019) demonstrated a reduced growth rate after exposing glass fish (*Ambassis dussumieri*) to microplastics for 92 days and a similar reduction in growth rate has been shown in the round goby (*Neogobius melanostomus*) (D’Avignon et al., 2023) and in Nile Tilapia (*Oreochromis niloticus*) (Mahmood et al., 2024) after an exposure period of 37 and 60 days, respectively. Interestingly, the reduction in growth rate in the Nile Tilapia was rather small even after 60 days of exposure (0.68% decrease). Shorter exposure periods have been shown to reduce growth rate for example in grass carp (*Ctenopharyngodon idella*) that assessed growth rate at seven, 14 and 21 days, and demonstrated an effect of microplastics on growth rate after day seven (Hao et al., 2023). Additionally, we assessed growth rate in mature, adult fish (90 dpf), whilst other studies that have shown a decrease in growth rate exposed fish from larval stages which are more sensitive to environmental stressors (Cattaneo et al., 2023). Furthermore, the size of plastic particles influences growth rate. Hao et al., (2023), showed that larger particles (15 μm) affected growth rate more than smaller ones (0.5 μm; the size of the particles used in the present study), whilst the smaller particles influenced other physiological traits more including stronger oxidative stress. Overall, there are clear intraspecies differences in responses to microplastics which vary with time scales, particle size and life stage.

Our study potentially provides evidence of behavioural compensation to retain growth rate, as fish exposed to microplastics had a higher probability of feeding and increased number of feeds which could offset reduction in digestive efficiency or nutrient assimilation caused by microplastic ingestion (Ferreira et al., 2016). Indeed, other studies have also shown an increased feeding rate during microplastic exposure (e.g., Uy and Johnson, 2022). This may be a compensatory response or, as it has been hypothesised, in some cases microplastics can induce changes in fish feeding behaviour through chemotaxis (Savoca et al., 2017). It is therefore possible that behavioural plasticity may mitigate short-term reductions in growth driven by microplastic exposure. Therefore, effects of microplastics on growth may well emerge in food-limited or food-variable environments, where compensation by *ad libitum* feeding is not possible, especially in combination with a reduction in food searching behaviour, such as exploration.

Behaviour is a crucial fitness component because it enables organisms to rapidly adjust their strategies in response to environmental stress (Smith and Blumstein, 2008). In the context of plastic pollution, behavioural traits appear particularly sensitive and are more readily altered compared to life-history traits (Mattsson et al., 2017; Pitt et al., 2018; Salerno et al., 2021). We found that fish exposed to microplastics were less bold, spent less time exploring novel environments, and altered their feeding behaviour, consistent with previous studies (Mattsson et al., 2017; Yin et al., 2019; Chen et al., 2022). Such shifts in behaviour may arise from neural disruption caused by microplastic accumulation in neural or sensory tissues. For example, earlier work in zebrafish has shown that microplastics can impair brain function (Chen et al., 2020; Yang et al., 2025). Alternatively, reduced boldness and exploratory behaviour in microplastic exposed fish may reflect a decline in physiological or energetic condition that is not captured by gross growth metrics. In fishes, risk-taking behaviours are often condition dependent and can respond to short-term energetic state, stress physiology or allocation trade-offs (Planas-Sitjà and Ioannou, 2025; Thelamon et al., 2025). Regardless of the underlying mechanism, our findings suggest that microplastics in fishes’ environment can generate behaviourally mediated fitness consequences. For example, reduced boldness may lead to poor reproductive success via mate choice as mismatched boldness between reproducing individuals is known to lower compatibility (e.g. causes differences in courtship frequency and responsiveness)(Ariyomo and Watt, 2012) and because females may avoid overly bold males (Herdegen-Radwan, 2019). Reductions in boldness also can reduce foraging efficiency (Magnhagen and Staffan, 2003), while a reduction in exploration may limit the ability to locate and exploit novel food resources (Conrad et al., 2011). Because microplastic exposure can increase feeding rate, likely to compensate for less nutritional intake, fish may also face elevated predation risk when allocating more time to foraging (McNamara and Houston, 1994; Pitcher, 1992). Additionally, microplastic ingestion can alter energy allocation away from key metabolic processes, or even induce an immune response (Limonta et al., 2019; Sadler et al., 2019), further contributing to potential fitness costs.

Despite the contrasting selective pressures experienced by the different selection lines, we detected no behavioural differences between them except in exploratory behaviour. This contrasts with earlier work on the same selection lines, which indicated consistent behavioural divergence among the lines (Roy et al., 2023; Sadler et al., 2024; Sbragaglia et al., 2019; Uusi-Heikkilä et al., 2015). However, the selection lines have undergone ten generations of recovery (i.e. no selection; van Dijk et al., 2024), during which behavioural differences may have eroded. Additionally, high variation among individuals and between selection lines may mask some behavioural differences, for example, Critchell and Hoogenboom, (2018) found an effect of microplastics on activity in one clutch of the spiny chromis (*Acanthochromis polyacanthus*), but not among the other two clutches tested.

Nevertheless, we do show that small-selected fish were less exploratory, consistent with previous work suggesting small-selected fish exhibit shyer behaviour (Monk et al., 2021; Sadler et al., 2024; Sbragaglia et al., 2019; Uusi-Heikkilä et al., 2015). A decrease in boldness in small-selected fish may correlate with decreased feeding and subsequent energetic shifts during selection (Uusi-Heikkilä et al., 2015). While a previous study demonstrated a difference in behaviour and growth rate between small- and large-selected fish in response to an aquatic pollutant, we showed that the selection-lines differed only in explorative behaviour in response to plastic pollution. This could be explained by the longer recovery time (i.e., ten generations; Sadler et al., 2024; van Dijk et al., 2024), and by the constraints typical with the large scale, long term selection experiments, replication of the selection lines are limited, where here we use two line replicates per size-selection, in other species (e.g., *Drosophila*) many more lines could be generated to understand repeatability of these evolutionary responses. It is noteworthy that while there was no significant interaction between growth rate and the selection lines, large-selected fish seemed to have somewhat lower growth rate than small- and random-selected fish, which is in line with previous studies (Uusi-Heikkilä et al. 2015,2024; Sadler et al. 2024; van Dijk et al. 2024). Nonetheless, these findings highlight that fisheries-induced evolution can shape how populations respond to emerging pollutants, with important implications for wild fish stocks exposed to both harvesting pressure and plastic contamination.

Our study demonstrates that short-term microplastic exposure primarily affects behavioural traits rather than growth in adult zebrafish, and that prior size-selective harvesting can mediate behavioural responses to pollution. These findings underscore the importance of integrating evolutionary history (e.g., previous fisheries selection), behaviour, and ecotoxicology when assessing the impacts of plastic pollution. Future studies should explore how multiple environmental stressors interact to shape demographic and evolutionary trajectories of fish populations in increasingly human-altered aquatic environments.

## Acknowledgements

We thank the JYU technical staff including Emma Pajunen, Mervi Koistinen, Noora Kinnunen, and all students that assisted with data collection.

