## Supplemental Information for "Microplastics influence size-selected zebrafish behaviour"

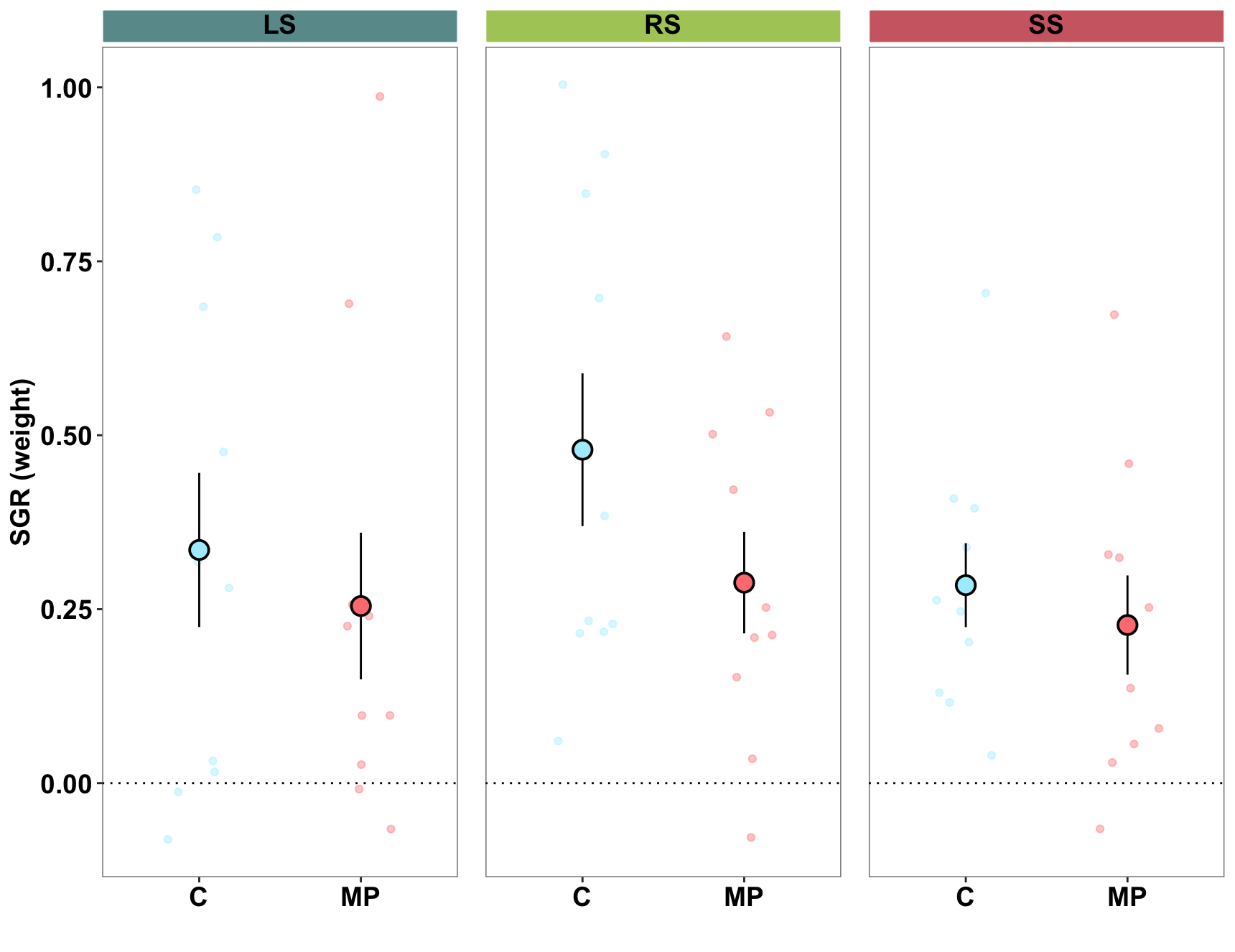

**Figure S1:** Differences in specific growth rate (SGR) in weight between control (C) and microplastic exposed fish (MP) across the selection lines. Large-selected (LS), random-selected (RS), and small-selected (SS). Large dots indicate mean ± CI. Small dots indicate individual replicates.

**Table S1**: Summary statistics of GLMMs/LMMs

| **Trait** | **Treatment** | **df** | **F/Z value** | **p value** | **Sig** |
| --- | --- | --- | --- | --- | --- |
| **Activity** | MP | 1 | 0.333 | 0.565 |  |
|  | Selection | 2 | 0.073 | 0.932 |  |
|  | MP: Selection | 2 | 0.032 | 0.968 |  |
| **Boldness** | MP | 1 | 7.503 | **0.007** | ****** |
|  | Selection | 2 | 0.593 | 0.606 |  |
|  | MP: Selection | 2 | 1.102 | 0.336 |  |
| **Exploration** | MP | 1 | 144.44 | **<0.001** | ******* |
| *GLMM* | Selection | 2 | 0.826 | 0.438 |  |
| *family = Poisson* | MP: Selection | 2 | 169.74 | **<0.001** | ******* |
| **Feeding latency** | MP | 1 | 0.362 | 0.549 |  |
|  | Selection | 2 | 0.302 | 0.759 |  |
|  | MP: Selection | 2 | 0.675 | 0.512 |  |
| **Feeding**  **probability** | MP | 1 | 5.491 | **0.019** | ******* |
| *GLMM*  *family = binomial* | Selection | 2 | 1.172 | 0.556 |  |
|  | MP: Selection | 2 | 0.837 | 0.658 |  |
| **Number of Feeds**  *GLMM*  *family = negative binomial* | MP | 1 | 12.12 | **<0.001** | ******* |
|  | Selection | 2 | 0.793 | 0.672 |  |
|  | MP: Selection | 2 | 0.062 | 0.969 |  |
| **Length (SGR)** | MP | 1 | 2.061 | 0.273 |  |
|  | Selection | 2 | 0.003 | 0.958 |  |
|  | MP: Selection | 2 | 0.518 | 0.503 |  |
| **Weight (SGR)** | MP | 1 | 0.003 | 0.958 |  |
|  | Selection | 2 | 2.061 | 0.273 |  |
|  | MP: Selection | 2 | 0.518 | 0.599 |  |
| **K Factor** | MP | 1 | 1.233 | 0.272 |  |
|  | Selection | 2 | 0.422 | 0.689 |  |
|  | MP: Selection | 2 | 0.912 | 0.408 |  |
